# The effect of heat on SARS-CoV-2 viability and RNA integrity as determined by plaque assay, virus culture and real-time RT-PCR

**DOI:** 10.1101/2020.11.02.365015

**Authors:** Jane Burton, Hannah Love, Kevin Richards, Christopher Burton, Sian Summers, James Pitman, Linda Easterbrook, Katherine Davies, Peter Spencer, Marian Killip, Patricia Cane, Christine Bruce, Allen D G Roberts

## Abstract

The effect of heat on SARS-CoV-2/England/2/2020 viability was assessed by plaque assay and virus culture. Heating to 56°C and 60°C for 15, 30 and 60 minutes led to a reduction in titre of between 2.1 and 4.9 log_10_ pfu/ml but complete inactivation was not observed. At 80°C plaques were observed after 15 and 30 minutes of heating, however after 60 minutes viable virus was only detected following virus culture. Heating to 80°C for 90 minutes and 95°C for 1 and 5 minutes resulted in no viable virus being detected. At 56°C and 60°C significant variability between replicates was observed and the titre often increased with heat-treatment time. Nucleic acids were extracted and tested by RT-PCR. Sensitivity of the RT-PCR was not compromised by heating to 56°C and 60°C. Heating to 80°C for 30 minutes or more and 95°C for 1 or 5 minutes however, resulted in an increase of at least three Ct values. This increase remained constant when different dilutions of virus underwent heat treatment. This indicates that high temperature heat inactivation of clinical samples prior to nucleic acid extraction could significantly affect the ability to detect virus in clinical samples from patients with lower viral loads by RT-PCR.

## 1. Introduction

The current pandemic of SARS-CoV-2 has led to an unprecedented global expansion in laboratory testing for the viral nucleic acids and antibodies against the virus. The severity of the disease, route of transmission and the lack of prophylaxis fulfils the criteria for the virus being handled as a BSL3/ACDP 3 agent, as are the related viruses SARS-CoV and MERS-CoV (1). Propagation of SARS-CoV-2 requires the use of BSL3/ACDP3 laboratories (2). In the UK, to provide the extremely high throughput required, the processing of clinical samples for diagnostic purposes has been derogated to BSL2/ACDP2 (3, 4). To protect laboratory staff conducting SARS-CoV-2 diagnostic testing and research outside of BSL3/ACDP3, effective methods of inactivating live virus present in clinical samples and other virus-infected material are essential.

Heating at 56°C for 30 minutes is commonly used to inactivate complement in serum samples for serological investigations. This has been shown to have no adverse effect on IgA and IgG ELISAs (5) and is also used to inactivate live virus present in clinical samples (6). This temperature has been found to be effective for many but not all viruses (7, 8). The effectiveness of heat on the inactivation of SARS-CoV-2 can be influenced by numerous factors including the type of sample, heat source, tube type and the length of time that the samples are heated (e.g. including or excluding the time taken for the samples to reach the target temperature). A recent comparison of available literature for the effect of heat on previously described coronaviruses (9) has shown considerable variation between and within studies. Overall, it concludes that for SARS-CoV and MERS-CoV, heating to 60°C for 30 min, 65°C for 15 min and 80°C for 1 min reduces virus infectivity by at least 4 log_10_. Previous studies using SARS-CoV have demonstrated that the efficacy of heat inactivation is reduced in samples with higher protein content (e.g. Foetal Calf serum (FCS), human serum or Bovine Serum Albumen) (10–12) For successful inactivation the British Standard for virus inactivation (13) recommends a 4 log_10_ or greater reduction in titre.

Complete inactivation of SARS-CoV-2 has been reported at 56°C after 45 minutes and 100 °C after 5 minutes (14), with an observed increase in titre between 15 and 30 minutes incubation at 56°C. Other studies suggest that SARS-CoV-2 can be inactivated in less than 30 minutes, 15 minutes and 3 minutes at 56°C, 65°C and 95°C respectively (15,16). Again, when testing at 37°C and 42°C, an increase in titre was observed between 15 and 30 minutes, and 30 and 60 minutes ref. For virus-spiked nasopharyngeal and human serum samples, greater than 5 log_10_ reductions in viral titres have been reported for heat treatments of 56°C for 30 minutes and 60°C for 60 minutes. In virus culture supernatant alone, virus titre reductions greater than 6 log_10_ for 95°C for 15 minutes have been demonstrated (17).

Most of the studies on the effect of heat on coronaviruses have used TCID_50_. This may be less sensitive than use of the plaque assay (10,17–19) to determine virus titre in heat-treated compared to untreated samples. None of the studies further subjected the samples to virus culture and RT-PCR to determine the presence of viable virus below the limit of detection of the TCIDso assay.

The aim of this study was to determine the effect of heat on the viability of SARS-CoV-2 in tissue culture medium using plaque assay to determine the titre followed by more sensitive detection of live virus by culture followed by RT-PCR. In addition, the effect of virus heat treatment on RT-PCR sensitivity was evaluated.

## 2. Materials and Methods

### 2.1. Virus

The virus stock used was a P3 working bank grown from SARS-CoV-2 Strain England 2, a clinical isolate taken during acute illness, propagated in Vero E6 cells. The stock was prepared by infecting 95% confluent Vero E6 cells with virus to an MOI of 0.005. Virus was harvested after 6 days. The titre was determined to be 7.0 x 10^5^ pfu/ml by plaque assay as described below.

### 2.2. Heat inactivation

#### 2.2.1. Test samples

For each heat treatment 3 x 1 ml volumes of virus in tissue culture medium (MEM + 4% FCS) were tested. Heat treatments of 56°C, 60°C and 80°C for 15, 30 and 60 min, 80°C for 90 min and 95°C for 1 and 5 min were tested.

#### 2.2.2. Heat treatment

Heating was carried out in an unlidded mini dry hot block. The temperature of the different wells in the block was validated before use using a Digitron thermal probe (2024T) inserted in a non-skirted Sarstedt tube (A2034) containing 1 ml of water. Heating was achieved by placing 1 ml volumes of virus in the hot block at the same time as an identical tube containing 1 ml of water and the thermal probe. The incubation time was started when the liquid in the control tube reached the required temperature (this usually took approximately 10 minutes). At the end of the incubation time tubes were placed on ice before the plaque assay was carried out. Untreated tubes were left on ice whilst the heating step was carried out.

### 2.3. Plaque assay

24 well flat-bottomed cell culture plates (Thermo Scientific) were seeded with 3.0 x 10^5^ /ml Vero E6 cells in 0.5 ml volumes of 2 x MEM medium containing Glutamax (2 x MEM Gibco) + 10% FCS (Sigma,) + 1 x (final concentration) NEAA (Gibco,) + 1 x antibiotic-antimycotic (Gibco) and incubated overnight at 37°C + 5% CO_2_. Before use the cells were checked for confluency. Each virus sample was tested in triplicate. Triplicate 10-fold dilutions up to 10^−6^ of each sample were made in a microtitre plate (Costar). The medium was removed from the 24 well plate and 100 μl of each dilution was added to the appropriate well. Plates were incubated at room temperature with occasional rocking for 1 hour, then 0.5 ml overlay (final concentration of 1.5% CMC (Sigma), 1 x MEM, 4% FCS and 1 x Anti-anti) added to each well. Plates were incubated for 72 hours at 37°C.

Plates were fixed for 1 hour with 4% formaldehyde in PBS then washed three times with water and allowed to dry. Plates were stained with 250 μl of 1% Crystal Violet (Sigma) for 10 −15 minutes, washed twice with water, dried and the number of plaques counted and recorded.

### 2.4 Virus detection by serial passage

500 μl of each heat-treated or untreated virus sample was added to a 12.5 cm^2^ flask of 80% confluent Vero E6 cells, allowed to adsorb for 60 minutes at room temperature then 2.5 ml of MEM + 4% FCS was added. Two negative control flasks to which 500 μl MEM + 4% FCS was added in place of virus, were set up in parallel. At the beginning and end of each passage, 140 μl samples of culture supernatant were added to AVL (560 μl; QIAamp viral RNA mini kit (Qiagen) in duplicate. After 10 minutes, 560 μl of 100% ethanol was added. Nucleic acids were extracted according to the manufacturer’s instructions and eluted in 60 μl AVE buffer. RT-PCR analysis was conducted as described in Section 2.6. After one week cells were observed for signs of cytopathic effect (cpe) by viewing under a low magnification microscope. Samples for which no cpe was observed were passaged using the above method up to 3 times to allow amplification of any low volumes of virus present in the sample. After the first passage a single positive (control) and negative flask were passaged on.

### 2.5 RNA integrity following heat treatment

SARS-CoV-2 virus was diluted to give 1 ml volumes of 7.0 x 10^4^, 7.0 x 10^2^ and 7.0 pfu/ml. One sample from each dilution was subjected to heat treatment as described in Section 2.2. Duplicate 140 μl samples were taken from each tube into AVL and RNA extracted as described above.

### 2.6 RT-PCR

Nucleic acids were stored frozen at −80°C until they were subjected to RT-PCR in suitable batches with initial (day 0) and final (day 7) samples from each passage in the same run. RT-PCR was carried out on an Applied Biosystems Fast 7500 PCR machine in standard run mode using the SARS-CoV E Sarbeco assay (20, 21) using MS2 as an internal extraction control (22). The master mix comprised E-gene F and R primers and TM-P (400 nM, 400 nM and 200 nM final concentration respectively), MS2 primers and TM probe (20 nM, 20 nM and 40 nM final concentration respectively), 4 x TaqMan® Fast Virus 1-Step Master Mix made up with molecular-grade nuclease free water (Ambion) to a final volume of 15 μl. The amount of template material added was 5 μl. Cycling conditions were 55°C for 10 min, followed by 94°C for 3 min then 45 cycles of 95°C for 15 s and 58°C for 30 s.

## 3. Results

### 3.1. Effect of heat on the viability of SARS-CoV-2

The effect of heat on SARS-CoV-2 virus viability as determined by plaque assay is shown in Figure 1 and Table 1. A 4 log_10_ or greater reduction in titre was consistently observed in all replicates after 56°C for 30 minutes, 80°C for 90 minutes and 95°C for 1 and 5 minutes. Significant variation in heat-treatment efficacy was observed between replicates. At 56°C and 60°C the plaque assay titre increased with longer periods of heat treatment (Figure 1). The sensitivity of the plaque assay was 3 pfu/ml.

**Figure 1.**
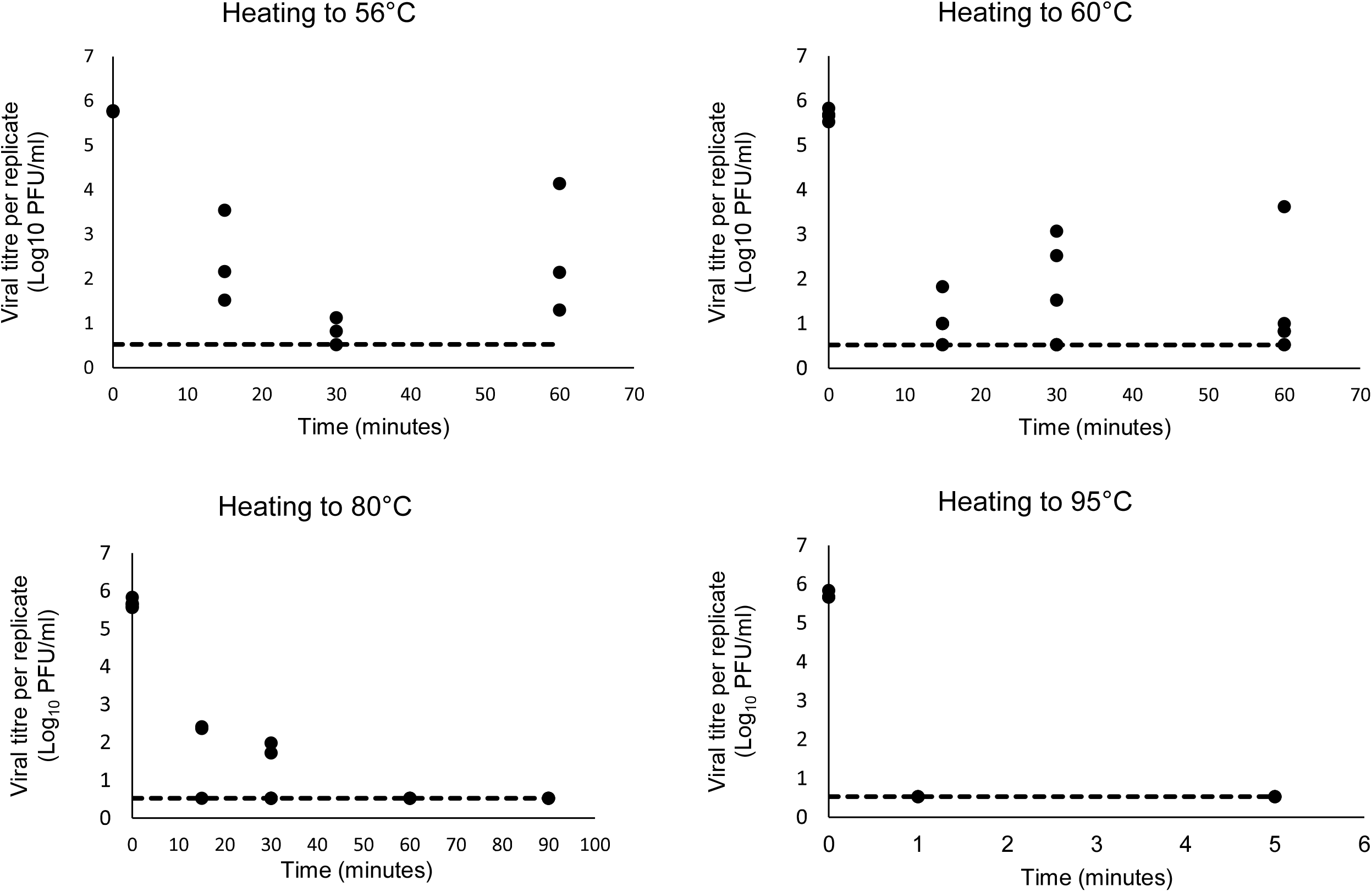
In each experiment the heat treatment was performed in triplicate on identical 1 ml volumes of SARS-CoV-2 England 2 virus in tissue culture medium. Each point represents the mean titre of three wells for each replicate at each temperature/time point (Log_10_ pfu/ml). Heat treatments at 60°C and 80°C were conducted twice (on separate days). The dotted line represents the limit of detection of the plaque assay.

**Table 1.** The effect of heat on the viability of SARS-CoV-2 England 2 virus assessed by plaque assay, observation of cytopathic effect (cpe)and RT-PCR

| Heat treatment |  | Log<sub>10</sub> titre reduction (pfu/ml) compared to untreated* |  |  | cpe observed / no. of replicates | Reduction in Ct value/ no. of replicates | Mean Ct value after heat-treatment | Standard deviation of Ct value |
| --- | --- | --- | --- | --- | --- | --- | --- | --- |
| Temperature (°C) | Incubation time (minutes) | Replicate 1 | Replicate 2 | Replicate 3 |  |  |  |  |
| Untreated | N/A | 0 (14 replicates) |  |  | 15/15 | 15/15 | 15.82 | 0.69 |
| 56 | 15 | 2.23 | 3.61 | 4.25 | 3/3 | 3/3 | 15.4 | 0.4 |
| 56 | 30 | 4.65 | – | 4.95 | 3/3 | 3/3 | 15.3 | 0.1 |
| 56 | 60 | 3.63 | 1.63 | 4.47 | 3/3 | 3/3 | 15.2 | 0.4 |
| 60 | 15 | 3.85 | – | – | 1/3 | 1/3 | 15.4 | 0.4 |
| 60 | 30 | 4.15 | 2.61 | 3.15 | 2/3 | 2/3 | 15.5 | 0.3 |
| 60 | 60 | 2.06 | – | 4.85 | 2/3 | 2/3 | 15.9 | 0.2 |
| 60 (2) | 15 | – | – | – | 2/3 | 2/3 | 16.2 | 0.4 |
| 60 (2) | 30 | 5.15 | – | – | 1/3 | 1/3 | 16.8 | 0.4 |
| 60 (2) | 60 | – | – | 4.85 | 3/3 | 3/3 | 16.3 | 0.3 |
| 80 | 15 | 3.36 | – | 3.31 | 2/3 | 2/3 | 17.8 | 0.6 |
| 80 | 30 | – | 4.00 | – | 3/3 | 3/3 | 20.2 | 0.9 |
| 80 (2) | 30 | 3.59 | – | – | 1/3 | 1/3 | 20.5 | 0.2 |
| 80 (2) | 60 | – | – | – | 1/3 | 1/3 | 25.5 | 2.5 |
| 80 (2) | 90 | – | – | – | 0/3 | 0/3 | 27.7 | 2.6 |
| 95 | 1 | – | – | – | 0/3 | 0/3 | 21.6 | 1.6 |
| 95 | 5 | – | – | – | 0/3 | 0/3 | 22.0 | 0.9 |

\* untreated control virus titre was 5.68 log<sub>10</sub> pfu/ml (St dev 0.1; mean 14 replicates); – = complete 5.68 log<sub>10</sub> titre reduction; no plaques were observed; (2) = duplicate experiment.

In all untreated and 56°C heated samples, cpe was observed and growth of virus was detected by SARS-CoV specific RT-PCR (as defined by a decrease in Ct value between nucleic acid samples extracted on day 0 and day 7) for each passage (Table 1). At 60°C, 3 of 6 replicates (from two separate experiments) showed virus growth (by cpe and RT-PCR) after 15 and 30 minutes and 5 of 6 after 60 minutes of heat-treatment. There was greater virus recovery after 60 minutes incubation at this temperature compared to shorter incubation times. At 80°C viable virus was recovered from 3 replicates where no plaques were observed. No virus was recovered or virus growth detected by RT-PCR from samples heated to 80°C for 90 minutes or 95°C for 1 or 5 minutes. All replicates showed 100% correlation between observation of cpe and detection of virus growth by RT-PCR.

### 3.2. The effect of heat on RNA integrity and RT-PCR assay sensitivity

Heat-treatment of virus to 56°C or 60°C had little effect on RNA integrity as measured by the sensitivity of RT-PCR (Table 1). Heating to 80°C for 30 minutes or more, however, resulted in an increase of at least three Ct values, equating to a log reduction in sensitivity of the RT-PCR. The Ct value further increased the longer the virus was heat-treated, with an increase often or more Ct values observed for samples held at 80°C for 60 or 90 minutes. Heating to 95°C for 1 or 5 minutes led to an increase in Ct of around 6. When different virus dilutions were heat-treated and tested by RT-PCR, the increase in Ct value observed after virus had been heated remained (Table 2) and as expected, greater variability between Ct values was observed when the initial RNA concentration was lower. After 60 minutes heating at 80°C the Ct value increased (compared to control virus that had not been heat-treated) by 10 and 14 for viruses at 7.0 x 10^4^ and 7.0 x 10^2^ pfu/ml respectively, whilst the virus at 7.0 x 10^1^ pfu/ml was rendered undetectable by RT-PCR.

**Table 2.** Ct values of SARS-CoV-2 RNA extracted from heat treated virus.

| Heat treatment |  | Ct Value |  |  |
| --- | --- | --- | --- | --- |
| Temperature (°C) | Time (minutes) | $7.0 \times 10^4$ (pfu/ml) | $7.0 \times 10^2$ (pfu/ml) | $7.0 \times 10^1$ (pfu/ml) |
| untreated | N/A | 17.2 | 23.2 | 30.2 |
| 56 | 60 | 18.6 | 24.3 | 30.5 |
| 80 | 15 | 18.1 | 24.8 | 28.8 |
| 80 | 30 | 21.0 | 29.6 | 29.6 |
| 80 | 60 | 28.1 | 37.5 | U |
| 80 | 90 | 25.8 | 35.4 | U |
| 95 | 1 | 18.8 | 26.6 | 32.5 |
| 95 | 5 | 21.7 | 29.4 | 36.2 |

Each Ct value is the mean from duplicate RNA extractions; N/A = Not applicable. U = Ct undetermined (RNA not detected)

## 4. Discussion

In this study a 4 log_10_ or greater reduction in virus titre was consistently observed in all replicates after 56°C for 30 minutes, 80°C for 90 minutes and 95°C for 1 and 5 minutes. Complete inactivation was observed after heating to 80°C for 90 minutes and 95°C for 1 and 5 minutes. As independent confirmation of our results, heat treatment at 56°C was also tested in a different PHE laboratory by TCIDso using a separate virus stock (concentration 7.0 log_10_ TCID_50_/ml). After 15, 30 and 60 minutes a 3.5, 5.3 and 4.3 log_10_TCID_50_/ml reduction in titre was observed. Taken together with our results, this is in agreement with results described by Pastorino *et al.* (18). It contradicts the findings of other studies where 56°C for 30 (15); (16)) or 45 minutes (14) or 80°C for 60 min (23) was shown to completely inactivate the virus. This variation is potentially due to differences in the sample volumes subjected to heat treatment. In this study 1ml volumes were tested to allow triplicate plaque assays and virus culture to be carried out. For high-throughput processing the volume of clinical sample required can be 600 μl. The other studies quoted tested volumes of 500μl or less. Another source of variation is the method of virus titration. This study used plaque assays (the gold standard for quantifying replication-competent lytic virions as plaque-forming units (24)), serial passage in cell culture and RT-PCR in combination to demonstrate the presence of viable virus below the limit of detection of the plaque assay and the effect of heat treatment on RNA integrity. These methodologies in combination are more sensitive than using titration alone. Another consideration is the potential variation in temperature that can occur when using a hot block. Although the hot block in this study was carefully monitored there have been reports that the use of hot blocks can result in unequal heating, hotspots, or spikes in temperature(25).

Interestingly, in this study at both 56°C and 60°C the number of plaques observed increased with longer treatment times. This is in agreement with findings using SARS-CoV (19) and has also been reported for SARS-CoV-2 at 56 °C (14) and at 37°C and 42°C (16). One possible explanation for this phenomenon may be the formation and subsequent dissociation of virus aggregates in response to heat (19). Virus aggregates would produce individual plaques in cell culture identical to those observed for individual virions, resulting in an underestimate of the true number of infectious virus particles in aggregated samples. True inactivation of disaggregated virions may occur after longer incubation times or at higher temperatures.

The reduction of virus RNA integrity observed in the RT-PCR assays correlated with the increased temperature and length of heat treatment. Integrity was not significantly affected by heating to 56°C and 60°C. This is in agreement with some studies (17, 26) but not others (27, 28). In this study we found that heating to 80°C for 30 minutes or more led to an increase in Ct value and therefore a reduction in RT-PCR sensitivity that could impact upon clinical diagnosis. The less notable increase in Ct value observed when virus was heated to 95°C in this study could be attributed to the shorter heating time. Further work needs to be done before in-depth conclusions can be drawn, however, preserving RNA integrity should be a consideration fordownstream processing requiring high sensitivity such as clinical diagnostic RT-PCRs. Heat treatment at lower temperatures combined with chemical inactivation, short duration high-temperature heat treatments, or chemical inactivation alone may be more appropriate to preserve RNA integrity and optimise PCR detection of SARS-CoV-2 RNA from low titre clinical samples.

In this study the effect of heat was tested on virus-infected tissue culture supernatant. It is likely that in some cases heat treatment would be even more variable in clinical samples although this has not always been reported (29). Our results show significant variation in the effectiveness of heat treatment for inactivation of SARS-CoV-2. This emphasises the importance of local validation of inactivation methods and the need for consistency in inactivation protocols to ensure sufficient reduction in virus titre for processing of clinical samples and research material in BSL2/ACDP2 facilities.

This research did not receive any specific grant from funding agencies in the public, commercial or not-for-profit sectors.

Declarations of competing interest of authors: None.

## Notes

### Competing Interest Statement

The authors have declared no competing interest.

